# Microfluidic platforms for probing spontaneous functional recovery in hierarchically modular neuronal networks

**DOI:** 10.1101/2025.05.25.654991

**Authors:** Keita Watanabe, Hideaki Yamamoto, Takuma Sumi, Hakuba Murota, Hironobu Osaki, Kenshiro Kawamoto, Teppei Matsui, Yoshito Masamizu, Shigeo Sato, Ayumi Hirano-Iwata

**Affiliations:** Research Institute of Electrical Communication (RIEC), Tohoku University, Sendai, Japan; Graduate School of Engineering, Tohoku University, Sendai, Japan; Advanced Institute for Materials Research (WPI-AIMR), Tohoku University, Sendai, Japan; Graduate School of Brain Science, Doshisha University, Kyoto, Japan

## Abstract

Inherent capacity to flexibly reorganize after injury is a hallmark of brain networks, and recent studies suggest that functional consequences of focal damage are strongly influenced by the network’s non-random connectivity. Although many of these insights have been derived from animal models and computational simulations, experimental platforms that enable bottom-up investigations of the structure–function relationships underlying damage and recovery processes remain limited. In this study, we used polydimethylsiloxane microfluidic devices to construct hierarchically modular neuronal networks that mimic the architectural features of the mammalian cortex. Laser microdissection was employed to selectively sever intermodular connections, enabling controlled damage to either hub or peripheral connections. Damage to hub connections led to delayed recovery, requiring more than three days for correlations to re-emerge. In contrast, peripheral damage resulted in faster recovery. Experiments that induced repeated injury to neuronal networks further demonstrated that recovery primarily occurred through the formation of alternative pathways rather than restoration of the original connections. These findings highlight how topological features of neuronal networks shape their response to injury and subsequent reorganization, providing mechanistic insights into the intrinsic self-repair capacity of biological systems.

## 1 Introduction

Stroke and traumatic brain injury disrupt motor and cognitive functions, with severe damage often leading to irreversible loss of regional brain functionality.^1–4^ In contrast, depending on the extent and location of the injury, the function can sometimes partially recover, highlighting the brain’s capacity for reorganization.^5–7^ For example, in macaque monkeys, ibotenic acid-induced lesions in the primary motor cortex hand area caused motor impairments that recovered with rehabilitation, involving early compensation by the ventral premotor cortex and later by peri-lesional primary motor cortex, indicating time-dependent network reorganization.^8^ In addition, rodent studies have shown that lesions in the primary motor cortex impair sensory processing in remote regions, which again can recover through the loss and gradual compensation of inhibitory connections.^9^ The importance of damage location in the recovery process is underscored by observations that damage to the internal capsule, a hub pathway connecting the cortex to the spinal cord, corticospinal tract, a hub pathway connecting the motor cortex to the spinal cord, results in prolonged motor impairment compared to a stroke directly in the cortex.^10, 11^ At a broader scale, computational simulations using a neuronal mass model in structured brain networks derived from diffusion magnetic resonance imaging have shown that the impact of focal lesions depends on network topology, with damage in central hubs disrupting widespread functional connectivity across the brain, while lesions in peripheral regions induce more localized effects.^12^ Collectively, these studies indicate that recovery after brain injury involves complex and dynamic reorganization across local and large-scale networks.

Recently, in vitro neuronal cultures are increasingly recognized as versatile platforms for elucidating the cellular and network mechanisms underlying the spontaneous recovery after injury, owing to their precise controllability of the extracellular environment and the ease of observation and manipulation.^13, 14^ In particular, the integration of cultured neurons with bioengineering tools has gained significant attention for guiding neuronal growth and reproducibly constructing well-defined networks with living cells.^15–18^ Among these approaches, microfluidic devices have become widely used due to their ability to precisely define structures and their ease of use making them an excellent method for patterning neurons. This capability enables real-time monitoring of axonal degeneration and regeneration following injury, providing a robust platform for investigating the mechanisms of damage tolerance and self-repair in living neuronal networks.^19^

To date, several in-vitro injury models utilizing microfluidic devices have been established. These include physical injuries induced by vacuum aspiration or compression,^20, 21^ chemical injuries caused by application of neurotoxins,^22^ and optical injuries achieved through laser microdissection.^23, 24^ Such approaches have contributed to understanding the effects of localized damage on individual axons and on randomly connected neuronal networks. However, despite these advances, an experimental platform that enables systematic evaluation of interactions between network architecture and injury yet remains to be established, leaving behind a comprehensive understanding of structure-function relationships at the network level.

In this work, we used microfluidic devices to constrain the growth of primary cortical neurons in dissociated culture and fabricated living neuronal networks bearing hierarchically modular structure—a network structure that is evolutionarily conserved in animal nervous systems.^25^ Using laser microdissection, we applied focal damage to microchannels coupling the modules and investigated the changes in spontaneous and evoked neuronal activity by calcium imaging. Experiments show that damage induced in hub and peripheral links has profoundly different effects on activity and its spontaneous recovery. Intriguingly, the recovery process was highly dependent on the topological role of the damaged connections even in this simplified model of biological neuronal networks: while hub damage led to a slower recovery than peripheral damage, it ultimately resulted in a greater rebound in network synchronization. This restoration was not solely attributable to the re-establishment of original connections but was also supported by the emergence of alternative functional pathways, indicating a compensatory reorganization of network activity.

## 2 Results

### 2.1 Laser microdissection of engineered neuronal networks

To induce localized damage to cultured neuronal networks, we constructed a fully optical system that integrates a 355 nm, nanosecond-pulse laser into an inverted microscope (Figure 1a). The laser beam is directed through mirrors and a dichroic mirror and is focused onto nerve fiber bundles within the microchannels using a 20*×* objective lens. The structure of the polydimethylsiloxane (PDMS) microfluidic device used to fabricate engineered neuronal networks is illustrated in Figure 1b. The network features a hierarchical modular architecture, consisting of a 2 *×* 2 groups, each of which is further subdivided into a 2*×* 2 array of smaller subgroups.^26, 27^ Each subgroup is implemented as a 100 *×* 100 µm^2^ through-hole, which we refer to as *modules*. The through-holes are interconnected by microchannels measuring approximately 6.0 *±* 0.9 µm (*n* = 12 microchannels) in height and 4.5 *±* 0.4 µm (*n* = 12 microchannels) in width. These microchannels restrict soma from entering while permitting extension of multiple neurites from the two ends. Primary rat cortical neurons were cultured in the microfluidic device and used for recordings at designated days in vitro (DIV).

**Figure 1.**
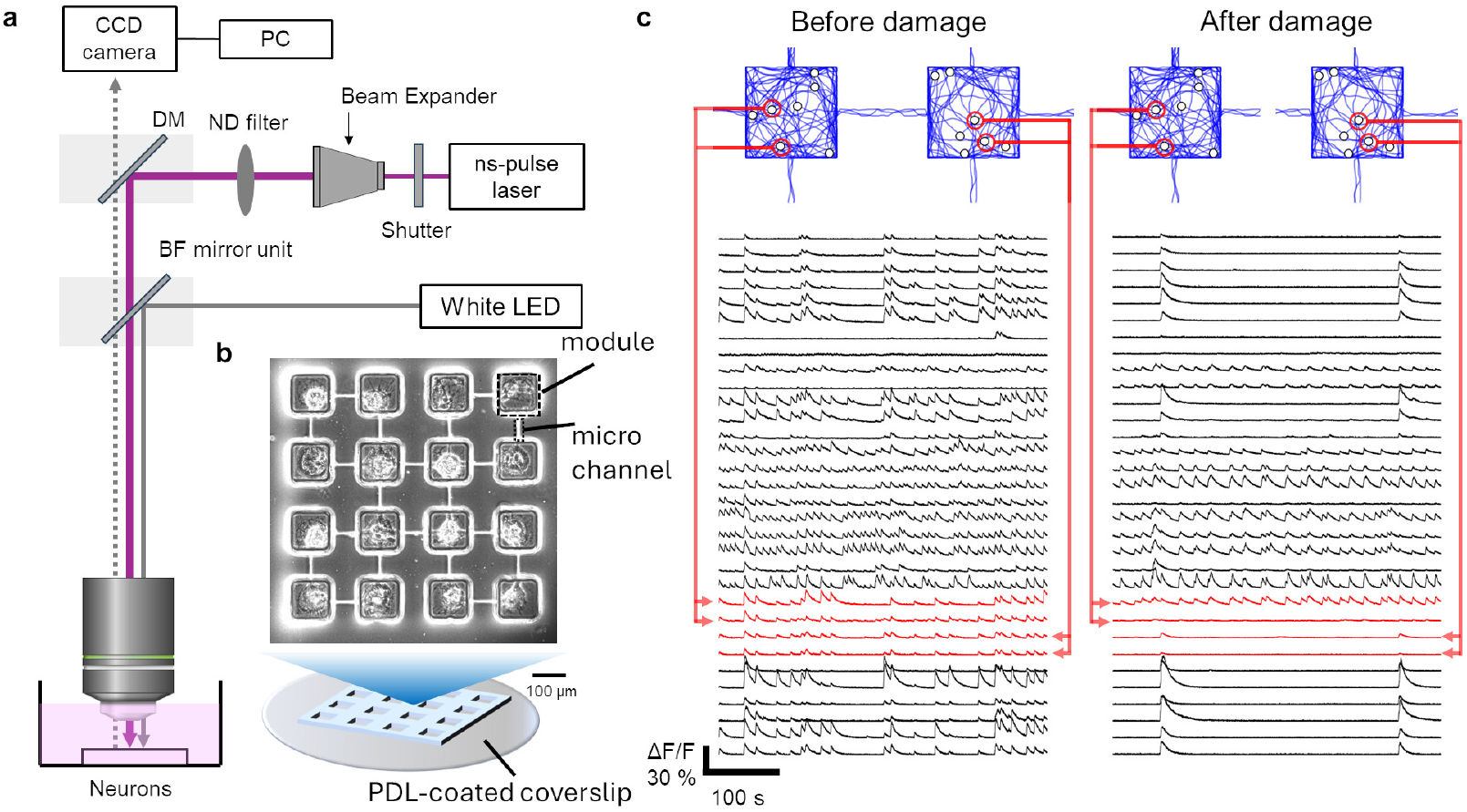
Overview of the experimental setup. a) Sketch of the laser microdissection system. b) Phase-contrast micrograph of an artificial neuronal network at DIV 10. A microfluidic device was attached to a PDL-coated coverslip and primary rat cortical neurons were seeded onto the device. c) Schematic illustrations of the injury experiment (upper panels) and representative fluorescence traces of GCaMP6s before and after injury (lower panels). Red traces represent neurons within the module corresponding to the damaged region. The traces are arranged from bottom to top according to the order of the ROI numbers shown in Figure. 3b. Schematics are not based on experimental data and are included for conceptual clarity.

Localized damage was induced in the network by irradiating a single laser pulse at the center of the microchannel region. This procedure selectively truncated neuronal processes within the microchannels while preserving the soma, thereby enabling targeted disruption of inter-modular connections without compromising the viability of nearby neurons (Figure 1c).

To assess neuronal viability following the damage, we compared the spontaneous activity of the neurons before and after laser irradiation. Prior to the damage, the entire network, including the neurons at the target site, exhibited a spatially and temporally complex activity that exhibited globally synchronized activity together with asynchronous activity (Figure 1c, left).^26–28^ Following the damage, globally synchronized activity diminished, and the firing rate of neurons at the damage site decreased. Nevertheless, these neurons continued to exhibit spontaneous activity, indicating that they remained viable (Figure 1c, right).

As a pilot experiment of laser microdissection, laser pulses were delivered to the two microchannels connecting the four modules on the bottom-right (Figure 2a, designated as group “B”) to the rest of the network. Experiments revealed that this intervention functionally isolated the group from the rest of the network, decreasing the degree of neuronal correlation between those in group B against those in the rest of the network (Figure 2b). Quantitatively, the correlation between neuron pairs involving group B (i.e., A–B, B–C, and B–D) decreased after the injury in 94.1% of the pairs (*n* = 2168 pairs), while the value for those not involving group B (i.e., A–D, C–D, and A–C) was 22.7% (*n* = 1883 pairs), indicating a selective loss of functional connectivity (Figure 2c). Not only the fraction of neuron pairs but also the difference in pair-wise correlation after damage was significantly larger for neuron pairs involving group B than in those that did not (Figure 2d). These findings indicate that laser-induced damage selectively impaired the functional connectivity of the targeted population, effectively isolating the fraction from the network. Interestingly, localized damage also induced widespread effects on the entire network. For example, the mean correlation in the damaged network decreased by 59.3 *±* 20.5% (*n* = 12 networks) immediately after the damage (Figure 2e). A reduction of 55.6 *±* 31.1% was also observed in the mean firing rate. In contrast, when the damage was not applied to the network, no significant changes in either the mean correlation or the mean firing rate was observed between the two recording sessions. In summary, laser microdissection, coupled with cell patterning based on microfluidic devices, allows targeted disruption of neuronal connectivity and widespread changes in network activity.

**Figure 2.**
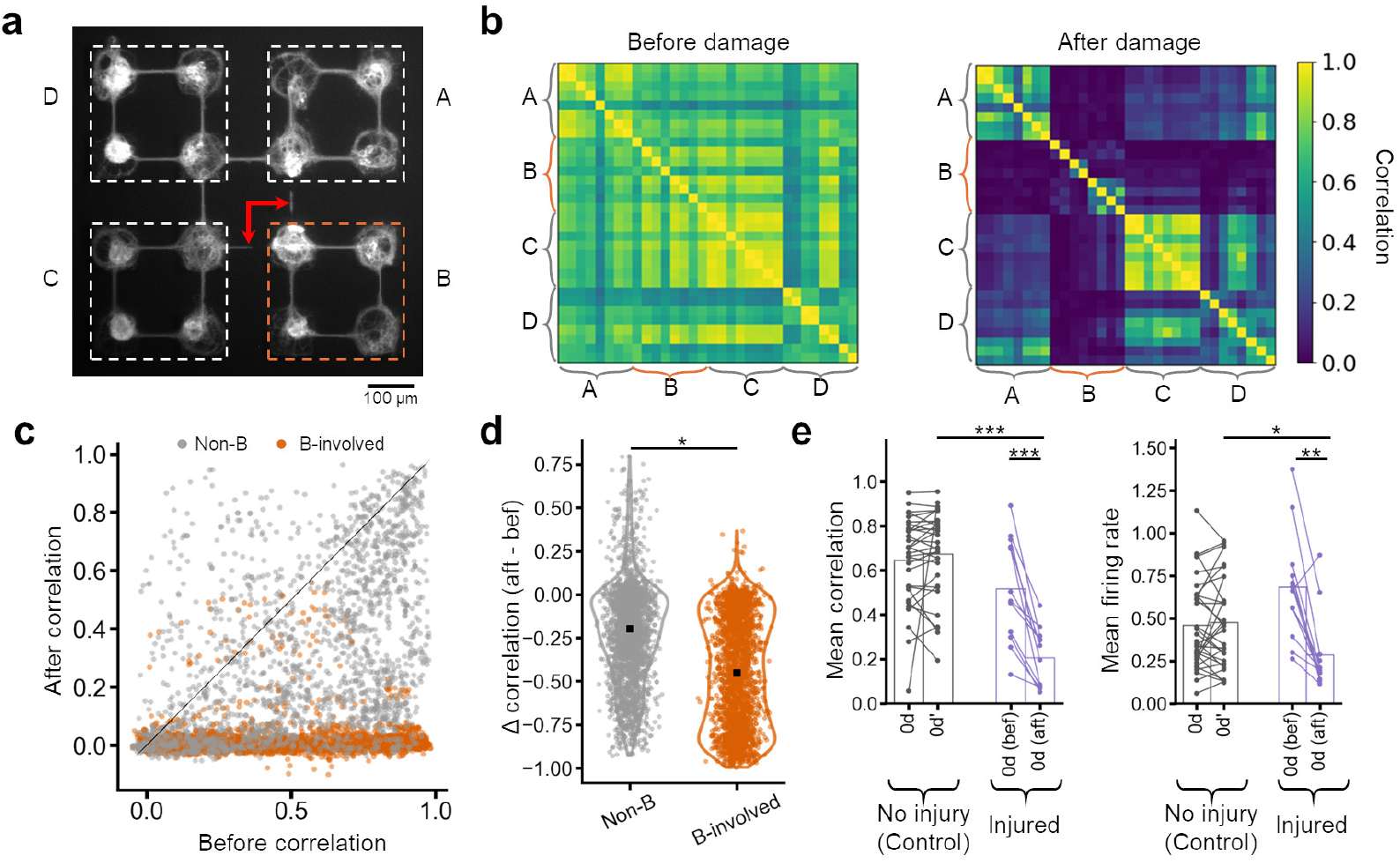
Effects of injury on artificial neuronal networks. a) Fluorescence image of the artificial neuronal network labeled with tdTomato under Synapsin-1 promoter. Arrows indicate the location of damage. b) Correlation coefficients between neurons calculated from estimated firing rates before (bef) and after (aft) the damage. c) Changes in correlation coefficients before and after the damage, grouped according to the spatial location of the neurons (*n* = 2304 pairs). Grey indicates pairs where both neurons belong to non-isolated modules; orange indicates pairs where one neuron belongs to an isolated module. d) Changes in correlation coefficients for individual neuron pairs (*n* = 2304 pairs). Black square dots represent the medians. Color coding corresponds to panel c). \**p* < 0.05 (Mann—Whitney U test). e) Comparison of neuronal activity between damaged (*n* = 12 networks) and undamaged (*n* = 31 networks) conditions. Left: Mean correlation coefficient across the whole network. Right: Mean firing rate across the whole network. Bars represent the means. \**p* < 0.05; \*\**p* < 0.01; \*\*\**p* < 0.001 (Wilcoxon signed-rank test for within-group comparisons; Mann–Whitney *U* test for between-group comparisons).

### 2.2 Effect of damage sites

To investigate how damage location influences the dynamics in a non-random network possessing hierarchically modular connectivity, we applied laser microdissection either to hub connections, which serve as central links, or to peripheral connections that connect terminal nodes. To assess both the impact of damage and the recovery process, spontaneous neuronal activity was recorded for 15 min at five time points: before the injury, immediately after the injury, and on days 1, 3, and 7 after the injury (Figure 3a).

**Figure 3.**
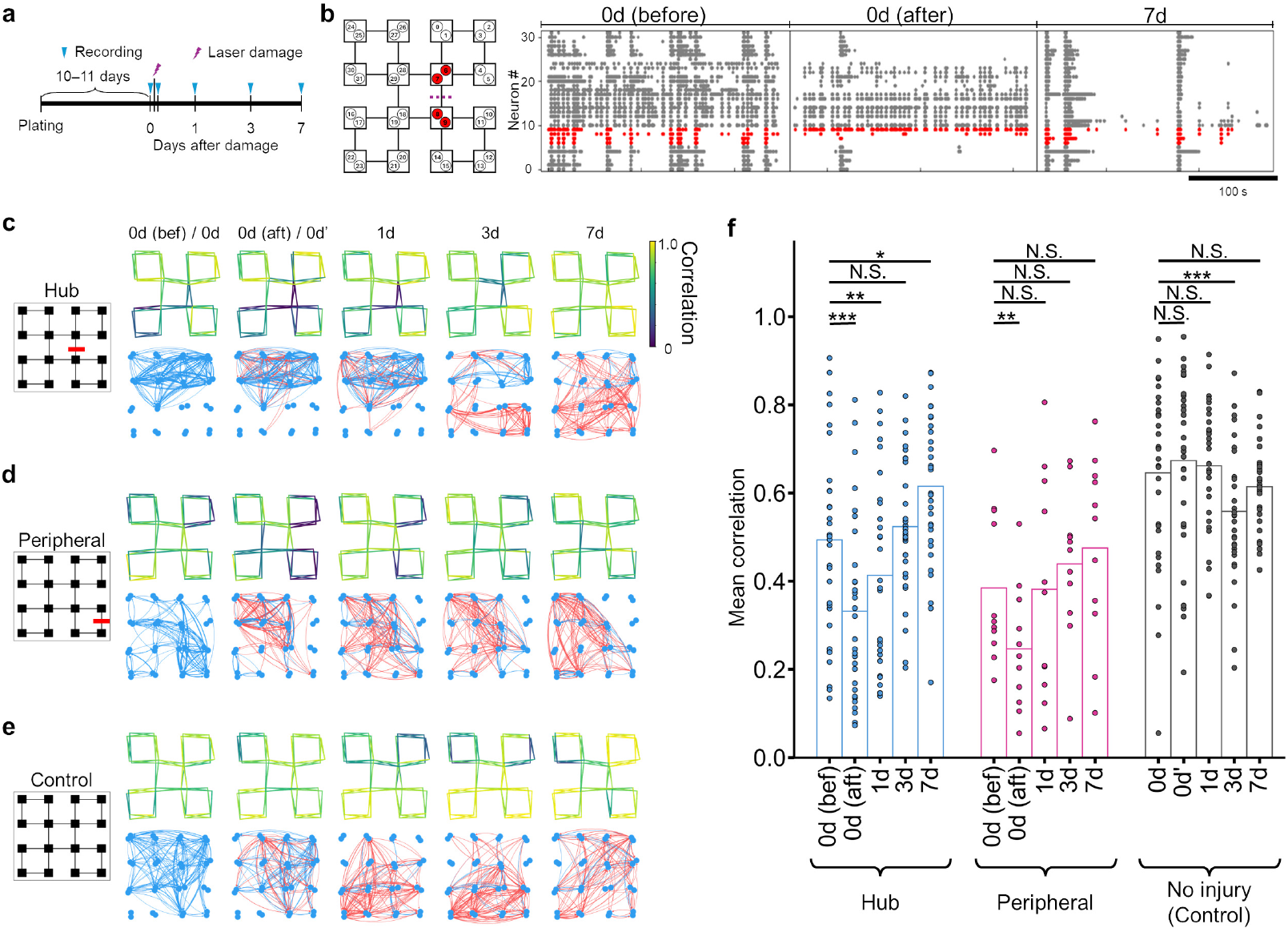
Hub and peripheral damage elicit distinct functional responses. a) Experimental protocol. Injury was induced by laser microdissection on DIV10–11. Spontaneous activity was recorded for 15 min at predefined time points: immediately before and after the damage, and 1, 3 and 7 days after the damage. b) Representative raster plots of neuronal activity recorded immediately before the injury (bef), immediately after the injury (aft), and 7 days after the damage (7d). The left panel illustrates the spatial layout of ROI, with neurons located near the injury site highlighted in red. The dashed purple line indicates the damaged hub connection. Neurons are ordered from bottom to top based on ascending ROI numbers. c–e) Representative maps of correlation-based connectivity and effective connectivity for networks with a damage in the (c) hub or (d) peripheral connections. A control network with no damage is presented in (e). The correlation map shows only structurally connected neuron pairs. Effective connections were estimated using transfer entropy, with values at each time point converted to *z*-scores. Only connections with *z* > 1 are visualized. Blue curves indicate connections that existed before the injury, while red curves indicate newly formed connections. Line thickness is scaled according to the *z*-scored value. f) Changes in the mean correlation across the entire network. Sample sizes: hub-damage (*n* = 32 networks), peripheral-damage (*n* = 11 networks), control (*n* = 31 networks). \**p* < 0.05; \*\**p* < 0.01; \*\*\**p* < 0.001; N.S., no significance (Wilcoxon signed-rank test with Bonferroni correction).

A representative progression of neuronal activity following damage to the hub connection is summarized in Figure 3b. Immediately after the injury, a reduction in firing rate and global synchrony was observed. By day 7, however, synchrony partially recovered. In particular, neurons nearby the damage location (shown in red) resumed to co-activate with other neurons. Irregular firing patterns and persistent decreases in activity were observed in some neurons, suggesting that recovery was heterogeneous across the network.

Figures 3c and d compare the changes in network activity following damage to either hub or peripheral connections. While a reduction in correlation coefficients was consistently observed immediately after damage, the timescale of recovery differed depending on the damage site (Figures 3c and d, upper panels). Evaluation of effective connectivity using transfer entropy further revealed that many of the original connections (blue) disappeared immediately after the injury, regardless of the injury site, and that new connections (red) emerge with development (Figures 3c and d, lower panels). In the absence of damage, the immediate reduction in neural correlation was not observed; however, a transition to new effective connectivity patterns was consistently observed (Figure 3e).

Quantitatively, focusing on the connections with a *z*-score *>* 1, 0.46 *±* 0.19 (*n* = 32 networks, hub) and 0.53 *±* 0.12 (*n* = 11 networks, peripheral) of the original connections were lost immediately after the injury. The fraction of original connections decreased through development, reaching 0.23 *±* 0.09 (hub) and 0.32 *±* 0.20 (peripheral) after seven days. Interestingly, no significant differences were observed between the hub and peripheral injury groups in the overall trajectory of these connectivity changes (Mann—Whitney *U* test), implying a common mechanism of network reorganization across injury locations.

Despite this large-scale reorganization, certain original connections still remained at day 7 after the injury. For example, stable edges remained within the four modules located in the upper right of Figure 3c and the upper left of Figure 3d. This suggests that functionally important “core structures” may persist even after injury, potentially serving as scaffolds for re-establishing global network connectivity.

To assess this trend, we compared changes in the mean correlation of the entire network under each condition (Figure 3f). In the case of hub connection damage, the mean correlation decreased by 35.7 *±* 19.0% after injury and required approximately three days to recover. Peripheral damage also caused a transient reduction by 33.3 *±* 25.4% but returned to the baseline level within one day. Notably, by day 7 after the injury, only the hub-damaged group exhibited a mean correlation exceeding the baseline level by 48.6 *±* 71.4%. These changes were not observed in the control group (without damage), suggesting that damage location plays a critical role in determining both the speed and extent of network recovery.

### 2.3 Mechanisms of spontaneous recovery

Finally, to elucidate the underlying mechanisms of this recovery, we used optogenetic tools to stimulate neurons in each module and analyzed changes in the propagation delay of the induced activity. The experiments were performed immediately after recording spontaneous activity on each measurement day (Figure 4a). To perform optogenetic stimulation, neurons were transfected with the red-shifted channelrhodopsin ChrimsonR at DIV 4 using viral vectors. Optogenetic stimulation was applied using patterned red light controlled by a digital mirror device, with the illumination area defined as a rectangle surrounding the neurons in each module. The stimulation sequence was set based on our previous study,^27^ with a duty cycle of 5%, a repetition frequency of 10 Hz, and an exposure duration of 1 s (Figure 4b). The stimulation across all 16 modules was repeated five times per experiment.

**Figure 4.**
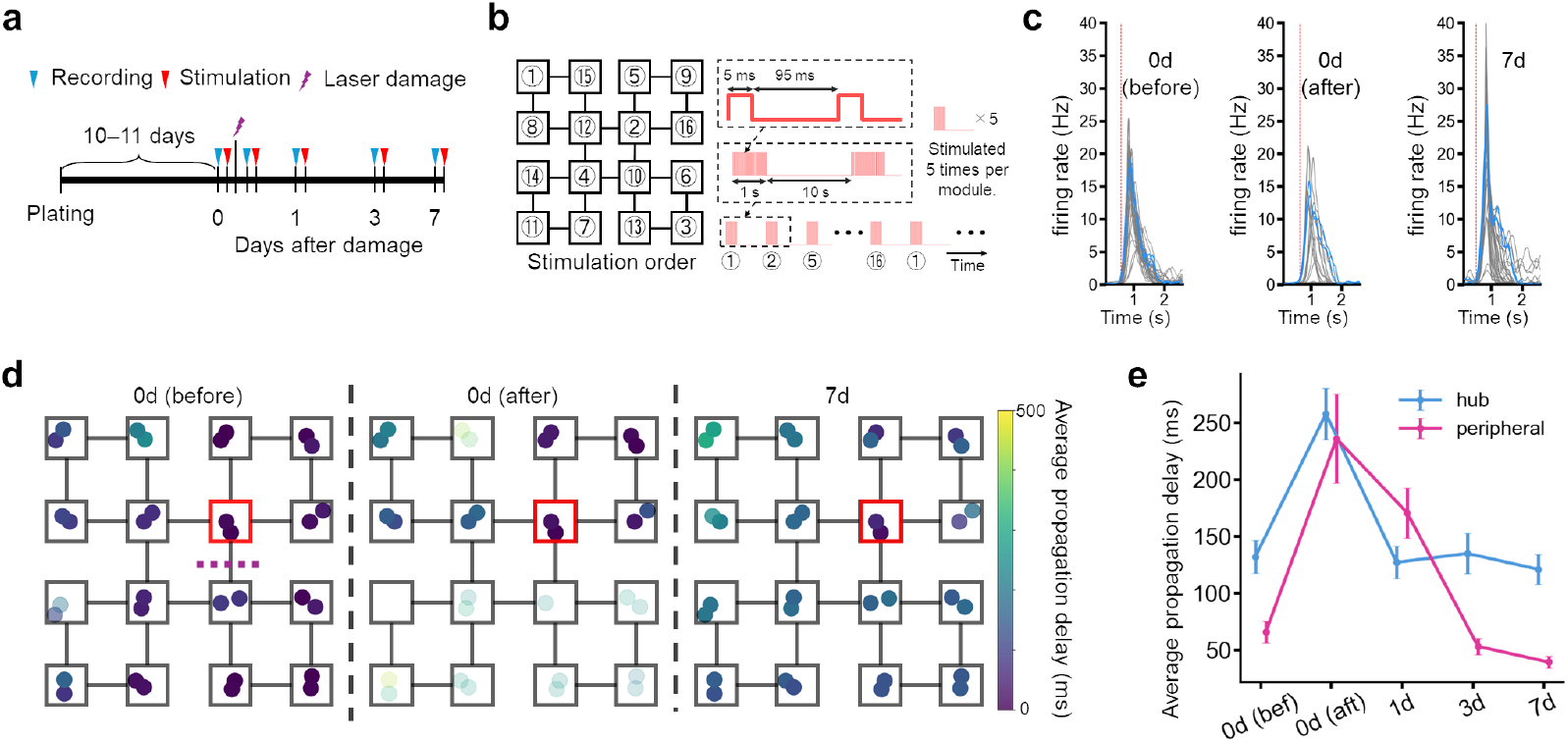
Propagation delay transiently increases following damage but recovers over time. a) Experimental protocol. After each recording session, the network was optogenetically stimulated to assess evoked responses. Other experimental conditions were identical to those described in Figure 3. b) Schematic description of the stimulation pattern. Each stimulus consisted of 5 ms light pulses delivered at 10 Hz for 1 s, with a 10 s interval between stimuli. c) Representative responses to a stimulation in module 2 upon a hub connection damage, shown for three time points: before the damage (bef), immediately after the damage (aft), and 7 days after the damage (7d). Blue and grey traces indicate neurons within module 2 and those in other modules, respectively. The red vertical line marks the time point at which any neuron within the stimulated module first exceeded a firing rate of 1 Hz. d) Propagation delay map of a representative network. The dashed purple line indicates the damaged hub connection, while the red square indicate the stimulated module. Propagation delays were spatially mapped onto the somatic locations. Colors represent the average delay, and transparency the response probability. Darker (more opaque) color indicates higher response probability. e) Evolution of average propagation delay. A hub connection damage was applied between modules 2 and 10 (*n* = 7 networks), while a peripheral connection damage was applied between modules 3 and 6 (*n* = 6 networks). Stimulation was applied to one of the two modules neighboring the damaged connection for five times, and the time required for the evoked activity to propagate to the other module was averaged for each sample. Plots and error bars represent the mean and the standard error of the mean (SEM), respectively.

Following hub connection damage, neuron-to-neuron variability in the rise and peak times of the evoked activity increased, but these timings realigned after seven days (Figure 4c). Moreover, analysis of propagation delays revealed that, prior to damage, responses could spread rapidly across multiple modules, whereas such propagation was suppressed immediately after the damage (Figure 4d). In particular, propagation of activity through damaged connections was inhibited. By day 7 after the damage, both the extent and delay of propagation approached the baseline levels, suggesting a recovery of the network’s signal transmission capacity.

Next, we investigated the temporal changes in propagation delay around the damaged areas. Specifically, in the hub damage condition, modules 2 and 10 were stimulated and the connection between them was damaged, whereas in the peripheral damage condition, modules 3 and 6 were stimulated and the connections between them was damaged. The mean propagation delays were then evaluated for each condition (Figure 4e). In the hub damage condition, the propagation delay increased from 132 *±* 139 ms to 258 *±* 138 ms, and gradually returned to 121 *±* 135 ms by day 7. In the peripheral damage condition, the delay similarly increased from 65.6 *±* 74.5 ms to 234 *±* 146 ms, and returned to 39.2 *±* 50.1 ms by day 7. These results show that signal propagation across microchannels restores spontaneously irrespective of the damage sites. Although propagation delays recovered over time, it remained unclear whether this reflected restoration of the original damaged pathway or the recruitment of alternative paths through other modules. To address this, we conducted a re-lesion experiment in which the same hub connection was damaged again seven days after the initial damage (Figure 5, *n* = 14 networks). As a control, a separate group received a first hub damage on day seven (*n* = 10 networks).

**Figure 5.**
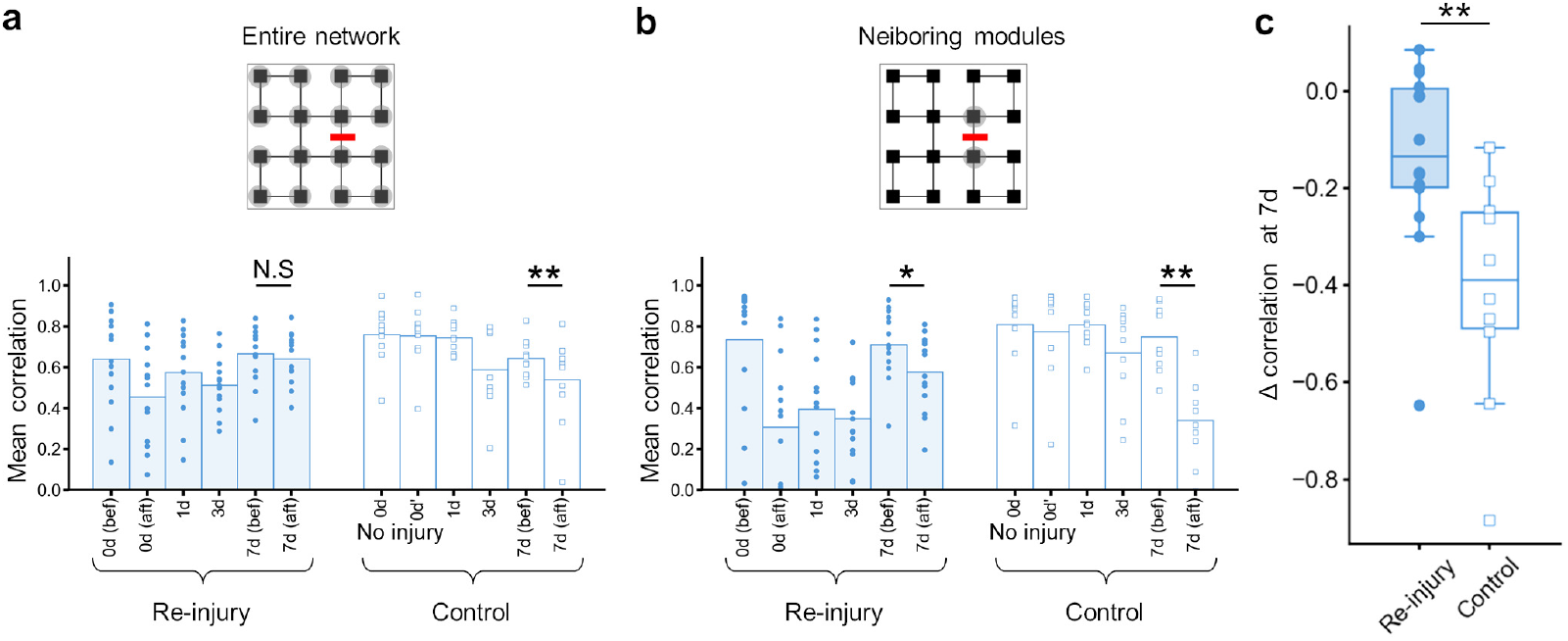
Prior injury increases the susceptibility of neuronal networks to subsequent perturbations. Sample sizes: re-injury (*n* = 14 networks) and control (*n* = 10 networks). a) Mean correlation coefficients across the network. \*\**p* < 0.01; N.S., no significance (Wilcoxon signed-rank test). b) Mean correlation coefficients and firing rates around the damaged region. Top: schematic of damage location (highlighted area indicates region of interest); bottom: mean correlation coefficient; bottom: mean firing rate.\**p* < 0.05; \*\**p* < 0.01; N.S., no significance (Wilcoxon signed-rank test). c) Change in correlation coefficients around the damaged region. \*\**p* < 0.01 (Mann–Whitney *U* test).

As was presented in Figure 3f, damaged networks exhibited an immediate reduction in correlation and mean firing rate following initial damage, with both metrics recovering by day 7. Interestingly, following the second lesion at day 7, networks did not show any significant change in the mean correlation. In contrast, networks subjected to the first damage on the same DIV showed a decrease in mean correlation by 18.1*±* 29.2%, confirming that the difference was not due to the timing of the initial damage (Figure 5a). This enhanced damage tolerance in the re-injury group was more pronounced when the metrics were evaluated for the neurons directly neighboring the targeted connections. Specifically, following the injury on day 7, the mean correlation decreased 16.4*±* 25.0% in networks that received a second injury, while the value was substantially greater (53.7 *±* 25.9%) for those that received their first injury (Figure 5b). Statistical comparison further revealed that the change in correlation coefficients following the damage on day 7 was significantly greater in the re-injury group (Figure 5c).

Summarizing, the results above point to two key mechanisms underlying spontaneous functional recovery in neuronal networks: (1) reconnection of original pathways, supported by the modest reduction in mean correlation and firing rate following the second injury, and (2) emergence of alternative routes that compensate for the lost connections, supported by the increased susceptibility in the re-injury groups. The increased susceptibility in the re-injury groups suggests that network reconfiguration, rather than exact restoration, plays a dominant role in recovery.

## 3 Discussion

We fabricated cultured neuronal networks with a hierarchically modular structure and selectively ablated specific inter-modular connections using focused laser pulses. By combining calcium imaging with optogenetic stimulation, we quantitatively tracked changes in spontaneous and evoked activity. Our findings revealed that (i) global network synchronization in the cultured neuronal networks, initially disrupted by the damage, gradually recovered over time, (ii) the rate and extent of functional recovery varied depending on whether the disrupted connections were hub or peripheral, and (iii) recovery was mediated not only by the reconnection in original pathways but also by the activation of alternative pathways. The third point suggests that recovery in neuronal networks involves functional compensation through reconstruction rather than a simple restoration of structure and function, providing complementary insights to the findings reported in animal and modeling studies.^29–32^ The approach presented herein enables bottom-up investigation of how physical damage impacts function in biological neuronal networks, providing insight into mechanisms that underlie robustness and functional recovery in complex networks. Microfluidic devices enable precise control of network connectivity in cultured neurons, uniquely positioning them as powerful tools to constructively investigate how structural features of neuronal networks influence their responses to perturbations. Various injury modalities—such as traction, compression, suction, laser ablation, and chemical treatments—have been shown to be compatible with microfluidic platforms to study axonal regeneration, axonal degeneration, neuroprotection, neuron–glia interactions, and other neural responses to injury.^20–22, 24, 33^ Our work extends these approaches to investigate injury responses in networks bearing a biologically plausible modular organization, clarifying the structural role of damaged regions and their influence on global network dynamics.

Previous in vitro studies have reported that localized injury to neuronal networks leads to a temporary reduction in firing rate and synchrony across the entire network, followed by partial recovery over the course of several hours to days.^23, 34^ In addition, a study combining experimental data with mathematical modeling has suggested that modular organization may contribute to redundancy in signal transmission and increase damage tolerance.^29^ In this study, we found that recovery of network activity differed depending on the location of the damage in the hierarchically modular network: disruption of hub connections led to delayed recovery and a prolonged desynchronization of the entire network. This indicates that the property of damaged regions critically influences the recovery process in structured networks of biological neurons. More broadly, such effect has also been observed in other complex networks, such as a power network and an Internet network (a supervisory control and data acquisition system), which are interdependent, and urban transportation networks.^35, 36^

Our findings are consistent with previous in vivo studies,^37–39^ which have shown that damage to central structures in brain networks significantly disrupts both structural and functional integration of the entire network. In particular, the widespread changes in network activity observed after damage to hub connections in our experiments closely match the functional impairments associated with injury to the internal capsule, a clinically important, highly centralized white matter structure. The internal capsule is a key structure in the central nervous system where the main conductive pathways connecting the cerebral cortex to the brain stem and spinal cord converge, and even small, localized lesions are known to affect diverse functional domains, including motor, sensory and cognitive functions.^40–42^ Furthermore, in the context of post-injury recovery, the formation of alternative pathways using residual structures is known to contribute substantially to functional recovery, alongside the regeneration of the originally damaged tracts, ^30–32^ further highlighting the strong parallels to our in vitro observations.

Damage to neuronal networks is common to many neurological disorders, and its understanding is essential for both basic and applied research. Our findings may not only contribute to the understanding of damage-induced functional impairment and the recovery process but may also lead to significant advances in the treatment of neurological disorders, the development of regenerative medicine, and the design of damage-resistant engineering systems.^43^

## 4 Experimental Section

### Laser microdissection

A system for inducing localized nerve damage was constructed by integrating a passive Q-switched solid-state laser (FTSS-355-50, CryLas; wavelength: 355 nm, pulse width: 1.1 ns; repetition frequency: 5 Hz) into an inverted microscope (IX83, Olympus). The laser beam was introduced into the optical path via a 355-nm compatible dichroic mirror, and then focused onto a bundle of nerve fibers inside a microfluidic chamber using a 20*×* water immersion objective (UMPLFLN 20×W, Olympus; numerical aperture NA = 0.5). To achieve efficient beam focusing, a UV-compatible beam expander (BE03-355-3X, Thorlabs) was integrated into the optical path to increase the beam diameter and collimate the beam. A neutral density filter (NDHN-U100, Sigma Koki) was used to adjust the laser power, with the final power at the sample surface set at 0.03 µW. A mechanical shutter (SSH-25RA, Sigma Koki) was used to isolate a single laser pulse by setting the open time to 0.2 s.

### Device fabrication

The microfluidic device was fabricated according to the previously reported protocol.^27, 44^ The master mold was fabricated by patterning two layers of SU-8 photoresist using photolithography. First, to form the first layer containing the module and microchannel structures, SU-8 3005 (Kayaku Advanced Materials) was spin-coated on a 2-inch silicon wafer at 3000 rpm. This was soft baked at 65 ^*°*^C for 1 minute followed by 95^*°*^C for 5 minutes, and then UV exposed through a photomask using a mask aligner (MJB4, Suss). This was followed by post-baking at 65^*°*^C for 1 min and 95 ^*°*^C for 3 min, developing with SU-8 developer, and washing with 2-propanol. For the second layer, SU-8 3050 was spin-coated at 1500 rpm, baked at 65 ^*°*^C for 1 min and 95 ^*°*^C for 30 min. A photomask was used to form only the module pattern on the second layer, followed by photolithography. The sample was post-baked at 65 ^*°*^C for 1 min and 95 ^*°*^C for 5 min. The development process was the same as for the first layer, and finally the mold was completely cured by baking at 60 ^*°*^C for 3 h.

The master mold was used to fabricate a microfluidic device by replica molding. Sylgard 184 (Dow Corning) was mixed with a 7.5:1 weight ratio of base and curing agent, degassed in a vacuum chamber, poured onto the master mold, and cured by heating at 70 ^*°*^C for 2 h. After curing, the PDMS was peeled from the mold, ultrasonically cleaned in 100% ethanol and deionized water, and then sterilized by exposure to ultraviolet light for over 30 min.

### Cell culture

All animal experiments and gene introduction procedures were performed under the approval of the Tohoku University Center for Laboratory Animal Research (2020AmA-001) and the Tohoku University Center for Gene Research (2019AmLMO-001). Primary rat cortical neurons were prepared from embryonic day 18 rat embryos following previously published protocols.^15, 44, 45^ Before cell seeding, a microfluidic device was attached to a poly-D-lysine coated coverslip and filled with plating medium [Minimum Essential Medium (MEM; 11095-080, Gibco) supplemented with 5% fetal bovine serum and 0.6% D-glucose]. To remove air bubbles from the microchannels and modules, the device was placed in a vacuum chamber under reduced pressure.

The medium was then replaced with fresh plating medium, and the coverslip was incubated in a humidified CO_2_ incubator (37 ^*°*^C, 5% CO_2_) for at least 3 h. Rat cortical neurons were seeded onto the coverslips at a density of 4.2 *×* 10^4^ cells/cm^2^, incubated for 3 h, and subsequently transferred to a culture dish containing 5 mL of Neurobasal Plus medium [Neurobasal Plus (A3582901, Gibco) supplemented with 2% B-27 Plus Supplement (A3582801, Gibco) and 0.25% GlutaMAXI (35050-061, Gibco)]. At DIV 4, half of the medium was removed and adeno-associated virus (AAV) vectors encoding the genetically encoded calcium indicator GCaMP6s (#100843-AAV1, Addgene) and the red-shifted channelrhodopsin ChrimsonR (#59171-AAV1, Addgene) were applied separately. On DIV 5, 2.5 mL of fresh Neurobasal Plus medium was added. On DIV 8, half of the culture medium was replaced with fresh Neurobasal Plus medium, and 50 µL of penicillin-streptomycin (P4333, Sigma) was added to prevent contamination.

### Calcium imaging

Calcium imaging recording was first performed on DIV 10 or 11, immediately before injury treatment. Subsequent recordings were performed under identical conditions immediately after the injury treatment and on days 1, 3, and 7 after the injury. For recording, the coverslip with micropatterned neurons was transferred to a glass-bottom dish (3960-03535, Iwaki) containing 1 mL of culture medium and 1 mL of fresh Neurobasal Plus medium. All subsequent culture and imaging were performed in the same dish. Culturing was continued until the final recording on day 7 after the injury. Following the recording on day 3 after the injury, half of the medium was replaced with fresh medium. Fluorescence intensities were acquired using an inverted microscope (IX83, Olympus) equipped with a 10× objective lens (UPLXAPO, Olympus; NA, 0.40), an LED light source (Lambda HPX, Sutter Instrument), a scientific CMOS camera (Zyla 4.2P, Andor), and an incubation chamber (Tokai Hit) with integrated temperature and CO_2_ control. During imaging, the CO_2_ concentration in the stage area was maintained at 5% using a CO_2_ controller, and the temperature was kept at 37^*°*^C. All recordings were performed using Solis software (Andor). Spontaneous neuronal activity was recorded at 20 frames per second, while responses to light stimulation were recorded at 100 frames per second.

### Optogenetic stimulation

Stimulation of the neuronal network was performed using patterned red light applied to neurons expressing the red-shifted channelrhodopsin ChrimsonR. The red light was emitted from a red LED light source (Solis 623-C, Thorlabs) and projected onto specific regions of the neuronal network using a digital micromirror device (DMD; Polygon400-G, Mightex). Stimulation pattern design and control were carried out using PolyScan2 software (Mightex).

### Data analysis

To extract neuronal activity from calcium imaging data, we manually selected 32 neurons from a single network (two neurons per module). Regions of interest (ROIs) were manually defined around each neuronal soma using Fiji software (NIH). The fluorescence intensity *F*(*t*) at each time point *t* was calculated as the mean pixel value within the ROI. For neurons with poorly defined somatic boundaries, a circular ROI with a diameter of approximately 15 µm, corresponding to the typical size of the soma. The relative fluorescence intensity (RFU, Δ*F/F*) for each cell was computed using the equation Δ*F/F* = (*F − F*_0_)*/F*_0_, where *F* represents the fluorescence intensity at time *t*, and *F*_0_ denotes the baseline intensity. The resulting RFU time series was converted into instantaneous firing rates *r*(*t*) using CASCADE, a deep learning-based algorithm for spike inference from calcium dynamics.^46^ The CASCADE model had been trained on a dataset consisting of simultaneous patch-clamp and calcium imaging recordings reported in the literature. To ensure compatibility with the trained model, the calcium signals obtained in this study were adjusted to match the same frame rate, noise level, and RFU amplitude (RFU = 1) as used during model training. Instantaneous firing rates inferred by CASCADE were used for subsequent analyses. This approach enabled the removal of signal artifacts caused by the rise and decay time constants inherent to the calcium indicator.

### Neuronal correlation

The functional correlation between neurons *i* and *j, CC_i j_*, was quantified using the Pearson correlation coefficient:

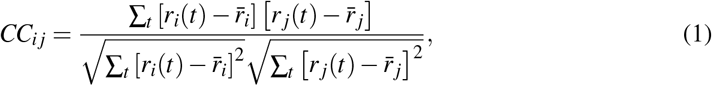

where, *r*_*i*_(*t*) and 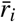 represent the instantaneous and time-averaged firing rate of neuron *i*, respectively. The mean correlation for a network was calculated by taking the mean of *CC*_*ij*_ across all neuron pairs excluding the diagonal elements.

### Propagation delay

To evaluate the propagation delay of neuronal activity between modules, we analyzed each cell’s response to optical stimulation using the following procedure. First, for each cell in the stimulated module, a firing event was detected when the instantaneous firing rate exceeded 1 Hz. The time point at which the firing rate first exceeded this threshold was defined as the reference time. The analysis time window was set to 1 s, spanning from 0.5 s before the onset of stimulation to 0.5 s before its offset. For cells in other modules, the response firing time was defined as the first point after the reference time at which the firing rate exceeded 1 Hz. The propagation delay time was then calculated as the time difference between the response firing time and the reference time.

### Effective connectivity

Causal interactions between neurons were quantified using Generalized Transfer Entropy (GTE).^47, 48^ Given spike trains of two neurons, *X* = *{x*_*t*_*}* and *Y* = *{y*_*t*_ *}*, the amount of information transferred from neuron *Y* to neuron *X* is evaluated as:

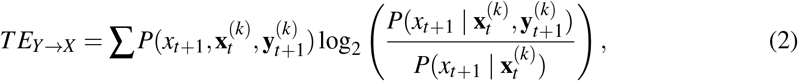

where *x*_*t*+1_ represents the state of neuron *X* at time *t* + 1, 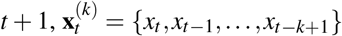, and 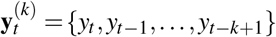 denote the past *k* states of neurons *X* and *Y*, respectively. In this study, the Markov order was set to *k* = 2.

Binary spike trains were obtained from 15 min spontaneous activity recordings with a time step of 50 ms. A spike was detected by applying a Schmitt trigger algorithm (upper threshold of 1 Hz and lower threshold of 0.5 Hz) to the instantaneous firing rate *r*(*t*) inferred using the CASCADE algorithm.^49, 50^ Once the upper threshold was crossed, the neuron was considered active until *r*(*t*) fell below the lower threshold.

### Statistical statement

All values in the analyzed data are presented as mean*±*SD or as means, unless otherwise noted. Sample size (*n*) is indicated in each section of the main text and in the figure captions. Statistical comparisons were performed using the two-sided Wilcoxon signed-rank test for paired (within-group) data and the two-sided Mann–Whitney *U* test for unpaired (between-group) data. For multiple group comparisons, *p*-values were adjusted using the Bonferroni correction.

## Acknowledgements

The authors thank Mr. Iori Morita and Mr. Nobuaki Momna at Tohoku University for the fabrication of the photomasks and for providing the code used to generate the figures. This work was partly supported by MEXT Grant-in-Aid for Transformative Research Areas (A) “Multicellular Neurobiocomputing” (24H02331, 24H02332, 24H02334, 24H02335), JSPS KAKENHI (22H03657, 22K19821, 22KK0177, 23H00251, 23H02805, 23H03489, 25H00447), the WISE Program for AI Electronics by Tohoku University, and the Cooperative Research Project Program of the RIEC, Tohoku University. This research was partly carried out at the Laboratory for Nanoelectronics and Spintronics, RIEC, Tohoku University.

## Conflict of Interest

The authors declare no conflict of interest.

## Data Availability Statement

The data that support the findings of this study are available from the corresponding author upon reasonable request.

## Notes

### Competing Interest Statement

The authors have declared no competing interest.

